# PaSD-qc: Quality control for single cell whole-genome sequencing data using power spectral density estimation

**DOI:** 10.1101/166637

**Authors:** Maxwell A. Sherman, Alison R. Barton, Michael Lodato, Carl Vitzthum, Michael E. Coulter, Christopher A. Walsh, Peter J. Park

## Abstract

Single cell whole-genome sequencing (scWGS) is providing novel insights into the nature of genetic heterogeneity in normal and diseased cells. However, scWGS introduces DNA amplification-related biases that can confound downstream analysis. Here we present a statistical method, with an accompanying package PaSD-qc (<u>P</u>ower <u>S</u>pectral <u>D</u>ensity-qc), that evaluates the quality of single cell libraries. It uses a modified power spectral density to assess amplification uniformity, amplicon size distribution, autocovariance, and inter-sample consistency as well as identifies aberrantly amplified chromosomes. We demonstrate the usefulness of this tool in evaluating scWGS protocols and in selecting high-quality libraries from low-coverage data for deep sequencing.

## Background

Whole-genome DNA sequencing of single cells (scWGS) has recently been made possible by the introduction of single cell amplification methods. Multiple displacement amplification (MDA) employs a highly processive polymerase which can synthesize new molecules (amplicons) of ~10-100 kb. High-quality MDA-derived data show that >90% of the human genome is amplified and 40-60% can be covered at >30X when the average depth is ~40-50X [1]. Copy number variations identified from low-coverage (<5X) MDA data have been used to elucidate tumor evolution [2] and to profile mosaic copy number variation [3]. With the decrease on cost of deep whole-genome sequencing, more recently, high-coverage (>30X) MDA data have allowed detection of somatic single nucleotide variants in the human brain [4]. Another protocol is Multiple annealing and looping based amplification cycles (MALBAC), which amplifies the genome in ~0.3-5 kb fragments and can cover ~50-90% of the human genome [5]. It has recently been proposed as a method for screening in-vitro fertilized embryos for genetic abnormalities prior to implantation [6, 7]. A third method based on DOP-PCR can amplify ~10% of the genome and is suitable for copy number variation detection but not single nucleotide variant detection [8].

All scWGS amplification methods induce biases and artifacts. These include non-uniform read depth that can appear as copy number aberrations, under and over amplification of entire chromosomes, uneven amplification of the two alleles, and correlation of features at the amplicon scale (e.g. ~10-100 kb for MDA) [9, 10], as well as single nucleotide and indel mutations and random ligation of fragments that are hard to distinguish from inversions. These biases fluctuate depending on the exact amplification protocol used and the state of the isolated cell (Figure 1A). For example, heat lysis during DNA extraction can increase the rate of artefactual C>T mutations compared to alkaline lysis [11], and cells in the G2/M phase amplify more uniformly than cells in the G1/G0 phase [12]. These biases in the data can affect the accuracy of variants detected in downstream analysis, and new protocols are frequently proposed claiming to mitigate these biases and provide superior variant detection [13, 14, 15]. It is thus important to characterize the biases computationally and assess the quality of single cell data.

**Figure 1:**
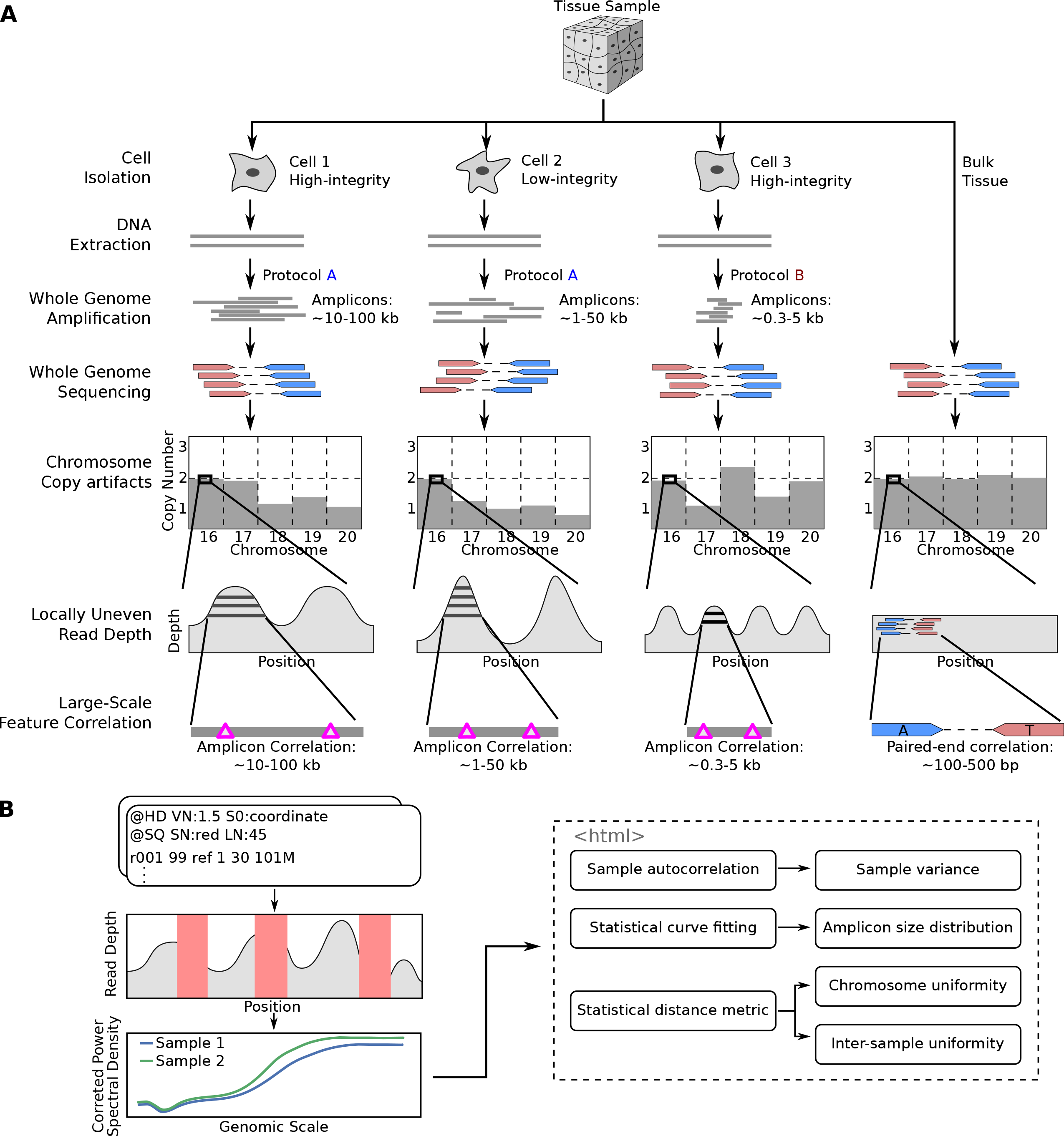
Overview of single cell whole-genome sequencing and sources of artifacts, and the PaSD-qc pipeline. **A**. Schematic overview of single-cell whole genome sequencing and the artifacts created by whole-genome amplification. The extent and patterns of the biases depend on the cell condition (high- or low-integrity) and on the scWGS protocol used (protocol A or protocol B). The pink triangles in the “Large-Scale Feature Correlation” represent genomic events (e.g., single nucleotide variants) which are spanned by a single amplicon and are thus correlated. The only correlation pattern present in bulk sequencing is due to paired-end sequencing, represented by positions marked “A” and “T” spanned by the mate pair. **B.** Schematic overview of the PaSD-qc pipeline. Read depth is extracted from bam files at uniquely mappable positions. Red rectangles represent regions where the true read depth is unknown due to low mappability, locus dropout, or sequencing bias. PaSD-qc uses a custom power spectral density estimation procedure to accurately estimate the correlation patterns in the data, and these patterns are then used to assess amplification properties and quality control measures. By default, the results are summarized in an interactive HTML report.

Despite the growing popularity of scWGS, few methods exist to perform this evaluation, and the few that do are almost exclusively concerned with estimating the uniformity of amplification. This itself is a non-trivial task because the true amplification process is masked by non-unique mappability, locus dropout due to amplification failure, or sampling bias during sequencing; additionally, read depth is highly correlated at positions spanned by the same amplicon. Current methods fail to account for these challenges. For example, several methods estimate read depth variance by binning reads [15, 16]. Such methods evaluate dispersion at a fixed genomic scale (the bin size), which fails to capture the correlation patterns of scWGS; resolving this requires re-binning at many scales, which is time-intensive and computationally expensive. More recently, an autocovariance (ACF) method has been proposed [10]. In theory, ACF is an appealing choice to capture the patterns in scWGS data because it measures correlations between observations within a dataset; however, in practice algorithms to estimate the ACF cannot easily be modified to account for regions of low mappability or locus dropout. Additionally, no standard implementations of these tools are available for incorporation into an scWGS pipeline.

Here, we introduce a suite of tools to comprehensively measure scWGS data quality, in a package called PaSD-qc (Power Spectral Density-qc, pronounced “passed-qc”). Using techniques from digital signal processing to estimate the power spectral density (PSD) of a sample and correct for observation gaps due to non-unique mappability, assembly gaps, and locus dropout without the need for binning, PaSD-qc provides a robust assessment of amplification uniformity at all genomic scales simultaneously. Because our estimation method accounts for the uneven spacing of the data while concurrently reducing background noise, the PSD can be leveraged to obtain more accurate estimates of variance and autocovariance than other methods; to identify chromosomes which may be copy-aberrant due to amplification failure in a principled way; and to compare quality across jointly analyzed samples even at very low coverage (<0.1X). Furthermore, our statistical method can estimate the full distribution of amplicon sizes in a sample, which has not previously been possible. PaSD-qc can easily be incorporated into existing pipelines and by default summarizes the quality and properties of each sample in an interactive HTML report. We use the tool to profile several different scWGS protocols, compare different samples from the same protocol, and select high-quality libraries from an initial set of low coverage (<1X) data for full-depth sequencing.

## Results

### Characterizing the spatial correlation structure induced by whole-genome amplification

Figure 1B provides an overview of PaSD-qc, and precise details of the algorithm are described in Methods. In brief, to mitigate issues of mappability, locus dropout, and sequencing bias, we extract read depth only at uniquely mappable positions covered by at least one read. The resulting signal is a time series (indexed by genomic position) with highly unevenly spaced observations. To infer the correlation patterns within this series, we apply the Lomb-Scargle algorithm [17, 18] to estimate the power spectral density (PSD) of the series. This method is one of the few which are capable of accurately analyzing correlation patterns of unevenly spaced time series data. We additionally apply a Welch correction [19] to minimize the noise of power spectral density estimation.

It is reasonable to ask (and was asked in [15]) whether the PSD is an appropriate approach given that it traditionally identifies periodic features when read depth is naturally an aperiodic signal. In fact, the PSD is mathematically equivalent to autocovariance, and for aperiodic signals, the PSD exactly estimates the variance of the generating process. See Supplemental Information (SI) for details. Thus, the smooth curves which result from our estimation method provide direct insight into the variance of scWGS amplification protocols at all length scales simultaneously.

Illustrative examples of a bulk sample, an MDA sample, and MALBAC sample are shown in Figure 2A. Below a genomic scale of ~1 kb, the samples show a characteristic pattern arising from paired-end sequencing. For a read pair with insert size *k* starting at positon *t*, there will be an increase in signal at *x_t_* and *x_t+k_* and a decrease in signal between the two reads. This results in periodicity at small genomic scales with the strongest periodicity at the mode of the insert size distribution (350 bp for the bulk sample shown). In fact, at small genomic scales, the PSD closely resembles the distribution of insert sizes in a sample (Figure S1). Above a genomic scale of ~1 kb, the bulk sample is virtually flat with low amplitude, indicating that, as expected, the coverage profile from bulk sequencing has low-variance and has no large-scale correlation structure. The slight increase in the PSD at scales >100 kb is an edge effect of the Welch correction. This edge effect is removed from scWGS PSDs by using an idealized bulk sample as a baseline (see Methods).

**Figure 2:**
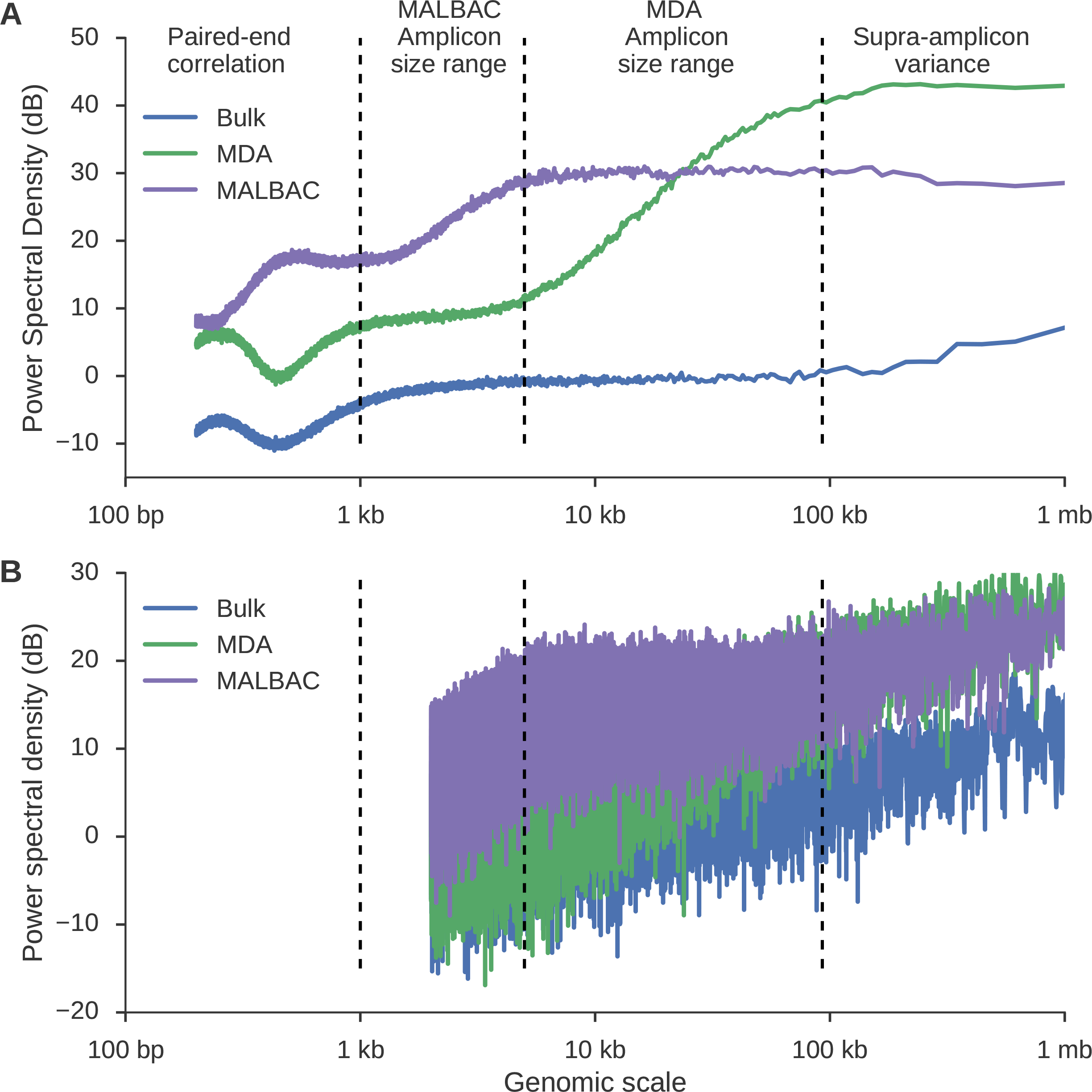
Using power spectral density to infer sample-specific amplification properties of scWGS data. **A**. PaSD-qc power spectral densities for a bulk sample (blue), MDA sample (green), and MALBAC sample (purple). The very low noise of the estimates allows amplification properties of the three samples to be inferred, including the paired-end insert size distributions (Figure S1), the range of amplicon sizes for MDA (~5-100 kb) and MALBAC (~1-5 kb), and the sub- and supra-amplicon variances of the two amplification protocols. Interestingly, whereas MDA has a higher supra-amplicon variance than MALBAC, its sub-amplicon variance is considerably lower. **B**. Power spectral density estimates using the algorithm from Leung et al, 2016 [13]. A similar algorithm is used in Zong et al, 2012 [5]. Background noise dominates the estimates making feature extraction infeasible. Resolution was limited to 2 kb because the data were binned into 1 kb bins as suggested per those algorithms.

The MDA and MALBAC curves have a more complex shape above the pair-end scale. To interpret these curves, consider an amplicon of length *h* starting at position *t*. The read depth signal *x_t_* will be correlated with *x_t+i_* for *i<h*. How often a correlation at length *i* is observed depends on the number of amplicons with length *h ≥ i*. If *i* is less than the smallest amplicon, then read depth *x_t_* and *x_t+i_* will almost always be correlated, resulting in small local variance and thus a lower amplitude PSD at sub-amplicon scales. For length *i* greater than the largest amplicon, *x_t_* and *x_t+i_* are necessarily independent, resulting in a higher amplitude PSD at supra-amplicon scales, reflecting the unevenness of the amplification. The PSD will smoothly transition from the sub- to supra-amplicon variances precisely following the cumulative distribution of amplicon sizes. These patterns are apparent in Figure 2A. The MDA curve rises from ~5-100 kb and the MALBAC curve rises from ~1-5 kb, consistent with expected amplicon sizes for these protocols. Additionally, the supra-amplicon variance of the MALBAC library is lower than the supra-amplicon variance of the MDA library while the opposite is true of the sub-amplicon variances, reflecting that MALBAC provides more consistent amplification at positions far apart but that MDA is locally more uniform since two positions close together are likely to be spanned by a single amplicon.

We are not the first to propose power spectral density estimation as a uniformity measure. However, prior estimation procedures [5, 13] require binning the data into 1 kb bins and do not take steps to reduce background noise. This results in an inferior PSD estimate where resolution is limited to a minimum genomic scale of 2 kb (since the Nyquist frequency is 5 × 10^−4^), and fine scale differences between samples are obscured by the high level of background noise (Figure 2B). Additionally, the PSD was criticized as lacking reproducibility since a Fourier transform may not be stable in regions of zero read depth and of low mappability [15]. As stated before, PaSD-qc corrects for these regions, resulting in highly reproducible estimates (Figure S2).

### Estimating the distribution of amplicon sizes in scWGS data

Since the dynamic region of the scWGS PSD curve reflects the cumulative distribution of amplicon sizes, this distribution can be estimated by fitting a properly scaled probability function to the PSD. The error function (erf) provides a particularly good fit (Figure 3A) and defines a density over the log amplicon sizes of the form *𝒩* (*μ,σ*^2^) (Figure 3B) where *μ* and *σ* are parameters estimated from the erf curve. In standard coordinates, the distributions are skewed with heavy tails extending into larger genomic ranges (Figure 3C). To confirm the accuracy of this estimation, we simulated an idealized amplification process on the p-arm of chromosome 3 using the amplicon size distribution estimated from the MDA curve as the generative model (see Methods). The resulting read depth signal was then analyzed using the PaSD-qc algorithm. Figure 3D shows the results of the simulation (red curve) along with the true estimate (green curve); Figure S3 shows the simulated curve for the MALBAC sample. In theory, any properly scaled and shifted sigmoidal cumulative distribution function can be used for this density estimation. We additionally tested the logistic distribution and gamma distribution as possible candidates, but found that the erf function produced the most consistent estimates (Figure S3).

**Figure 3:**
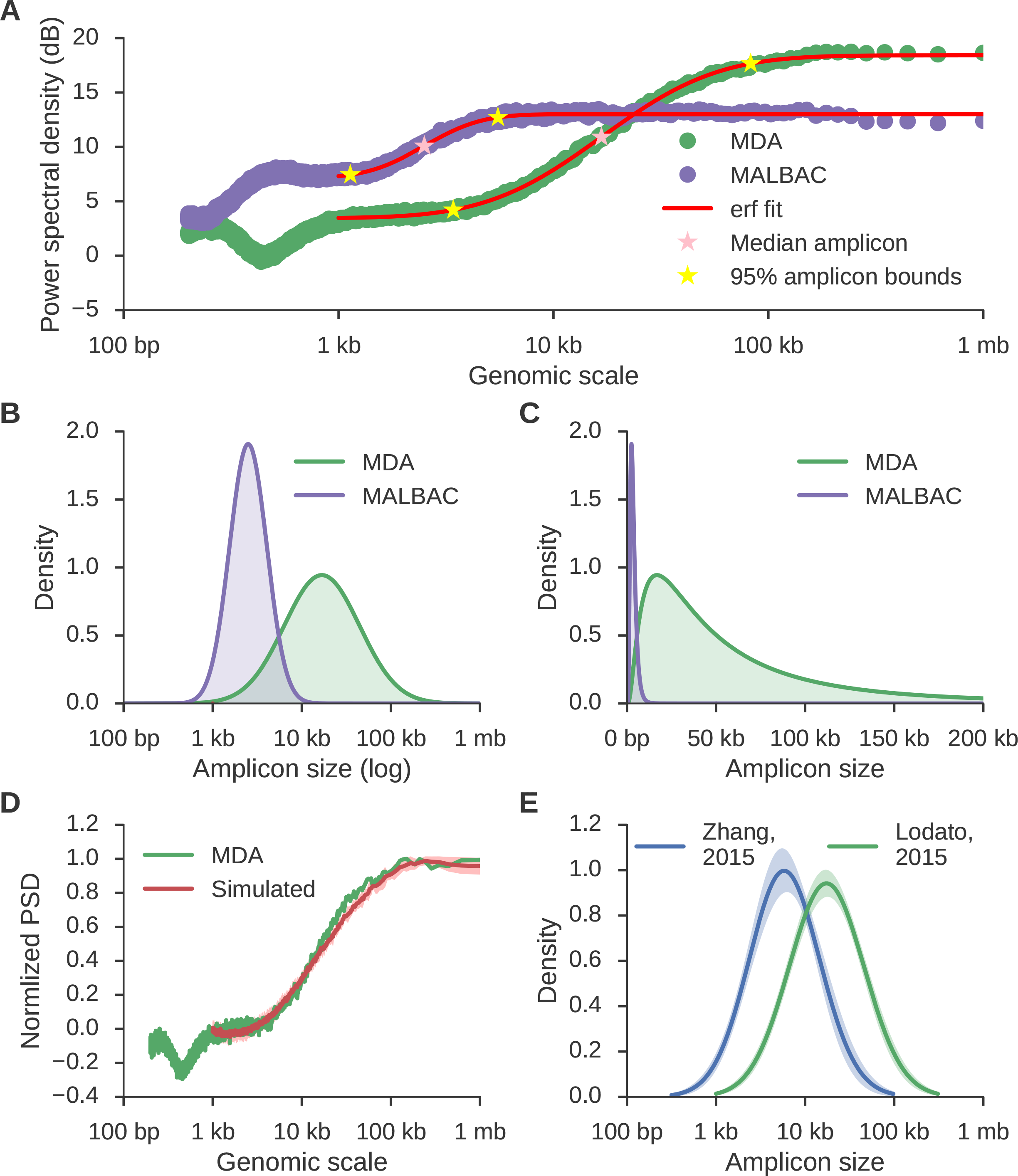
The distribution of amplicon sizes can be directly estimated from the power spectral density. **A**. MDA (green) and MALBAC (purple) curves as in Figure 2 along with the inferred error function (erf) fit of the dynamic region (red), the median amplicon size (pink stars), and 95% bounds on amplicon sizes (yellow stars). **B,C**. Distributions of inferred amplicon sizes in the MDA and MALBAC sample. Densities are normally distributed in a log scale (B), but highly skewed in standard coordinates (C). **D**. The average power spectral density (red) resulting from ten simulated amplification processes using the MDA density shown in C as the generative distribution. The shaded region represents the 95% confidence interval and the green curve corresponds to the original data. The MALBAC fit and fits using other distributions are shown in Figure S3. **E**. The average amplicon size distributions for 35 samples from Lodato et al, 2015 [4] (green) and 14 samples from Zhang et al, 2015 [20] (blue) reveal that different MDA protocols produce different amplicon size distributions, but a single protocol produces consistent amplicon size distributions across samples (shaded regions represent 95% confidence intervals around the average).

The median and percentiles of the amplicon sizes per sample are inferred using Monte Carlo simulation (see Methods). For the MDA sample, the median amplicon size is 16.7 kb and 95% of all amplicons fall between 3.3 and 89.0 kb in size; for the MALBAC sample, the median amplicon size is 2.5 kb and 95% of all amplicons fall between 1.1 and 5.6 kb in size (Figure 3C). We additionally profiled the amplicon distribution of 35 samples from [4] and 14 samples from [20] and found that while distributions are consistent between samples amplified with the same protocol, they are divergent between different protocols (Figure 3E); samples using the Qiagen REPLI-g Single Cell Kit with heat lysis [20] have a smaller median amplicon size than samples using Epicenter RepliPHI Phi-29 with alkaline lysis [4] (17.5 ± 1.3 kb vs. 5.9 ± 0.5 kb, p-value: 1.2e-9 by Kolmogorov-Smirnov test). We profiled 4 samples also profiled in that study and found that our estimated median amplicon size was consistent with their characteristic length scale estimate (Table S1). Although the characteristic length scale of correlation has been calculated before [10], no other method estimates the full distribution of amplicon sizes in scWGS data.

### Comparison to existing scWGS quality control metrics

The autocovariance function (ACF) of scWGS data has previously been proposed as a quality metric. While the ACF can be calculated directly from unevenly spaced time series data in theory, no computationally efficient algorithm exists to perform the estimation, and implementations are either time intensive, memory intensive, or both. Additionally, the statistical power at each lag varies and no theoretical results exist on the consistency of the unevenly spaced ACF estimator. However, it is possible and theoretically justified to calculate the ACF from the PSD (see SI). PaSD-qc implements an efficient algorithm based on this principle (see Methods).

To compare the performance of the PaSD-qc ACF against the directly calculated estimate, we analyzed all 16 single cell samples from the “1465” individual in [4] using both methods (Figure 4A). These samples were pair-end sequenced with an average insert size of 350 bp. The PaSD-qc ACF estimate consistently identifies the peak in correlation expected at this scale; the direct estimation fails to capture this feature. Additionally, the autocorrelation should oscillate around zero beyond the largest amplicon size. While this behavior is present in the PaSD-qc ACF, the direct estimation remains positive beyond 1 mb, a genomic scale far larger than the upper amplicon size limit of the Phi-29 polymerase used in MDA. This empirically demonstrates the potential inaccuracy of directly calculating the ACF from highly unevenly spaced observations and illustrates how PaSD-qc surmounts this limitation.

**Figure 4:**
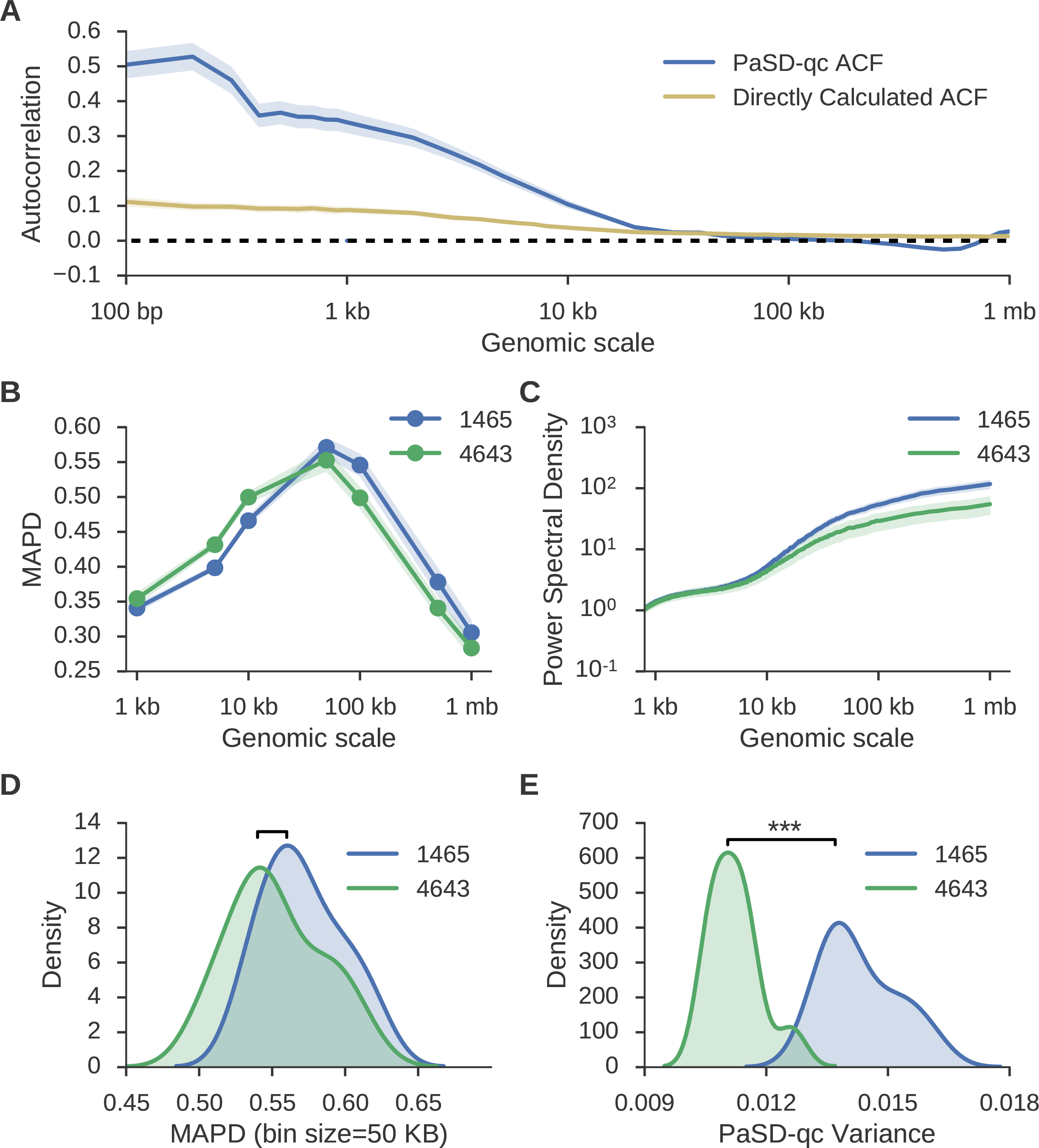
The PaSD-qc variance measure outperforms prior dispersion estimates. **A**. Average sample autocovariance with 95% confidence intervals for the 16 “1465” samples from Lodato et al, 2015 [4] as calculated by PaSD-qc (blue) and by direct estimation (gold). See text for a comparison. **B**. Average MAPD scores with 95% confidence intervals calculated for seven bin sizes ranging from 1 kb to 1 mb for 16 “1465” samples and 11 “4638” samples from Lodato et al, 2015. **C**. The average power spectral density with 95% confidence intervals for the same samples. **D**. Densities for the MAPD scores of the two sets of samples at 50 kb, the standard bin size at which the score is calculated. At this bin size, MAPD cannot distinguish behavior of the two sets of samples. **E**. Densities of PaSD-qc variance for the two sets of samples are significantly different.

Additionally, the ACF at lag zero (equivalently the integral of the PSD) provides an estimate of the overall variance. This dispersion estimate outperforms the other commonly used dispersion estimate, median absolute pairwise difference (MAPD) [16, 21]. MAPD is calculated by binning the read depth signal into fixed-width bins, calculating the normalized copy number in each bin, and taking the median of the pair-wise differences between all neighboring bins. We calculated MAPD scores at a range of bin sizes (Figure 4B) and the PaSD-qc PSD estimates (Figure 4C) for all “1465” and “4643” samples from [4]. Both reveal “1465” samples have higher supra-amplicon variance than “4643” samples. However, calculating MAPD even at a single bin size is computationally intensive; as such, it is usually calculated only for a single bin size, often 50 kb. At this scale, MAPD fails to distinguish a difference between the sets of samples (Figure 4D, p-value: 0.11 by Kolmogorov-Smirnov test). However, the PaSD-qc variance readily discriminates the two sets (Figure 4E, p-value: 1.7e-6 by Kolmogorov-Smirnov test).

### Identification of chromosomes with copy number altered due to aberrant amplification

The close relationship between a power spectral density estimate and a normal distribution [22] permits the calculation of a statistical distance measure, the symmetric Kullback-Leibler (KL) divergence, between two spectra (see Methods). For a given sample, PaSD-qc identifies chromosomes with aberrant amplification patterns by calculating the distance of each chromosome's PSD from the sample-average PSD. A chromosome is considered aberrant if it's KL-divergence is two standard deviations beyond the sample median across all chromosomes.

We demonstrate how this method can identify false-positive chromosomal copy alterations by analyzing the “1465” neurons from [4]. This set of samples is informative as it includes a high coverage bulk sample from the same tissue to establish a true copy profile. PaSD-qc identifies chromosomes 15-17 and 19-22 as inconsistently amplified in at least half of the samples (Figure 5A, Table S2); sex chromosomes are ignored in this analysis. Except for chromosome 15, each of these chromosomes is called as significantly copy-altered in at least 25% of samples (Figure 5B, Table S2) by the BICseq2 algorithm [23]. If a whole chromosome deletion is present in at least 25% of cells, that deletion should be apparent in bulk sequencing; however, bulk analysis of the tissue reveals all chromosomes to be copy neutral, indicating that the single cell copy alterations are artifacts. Different scWGS protocols show different patterns of aberrantly amplified chromosomes (Figure S4).

**Figure 5:**
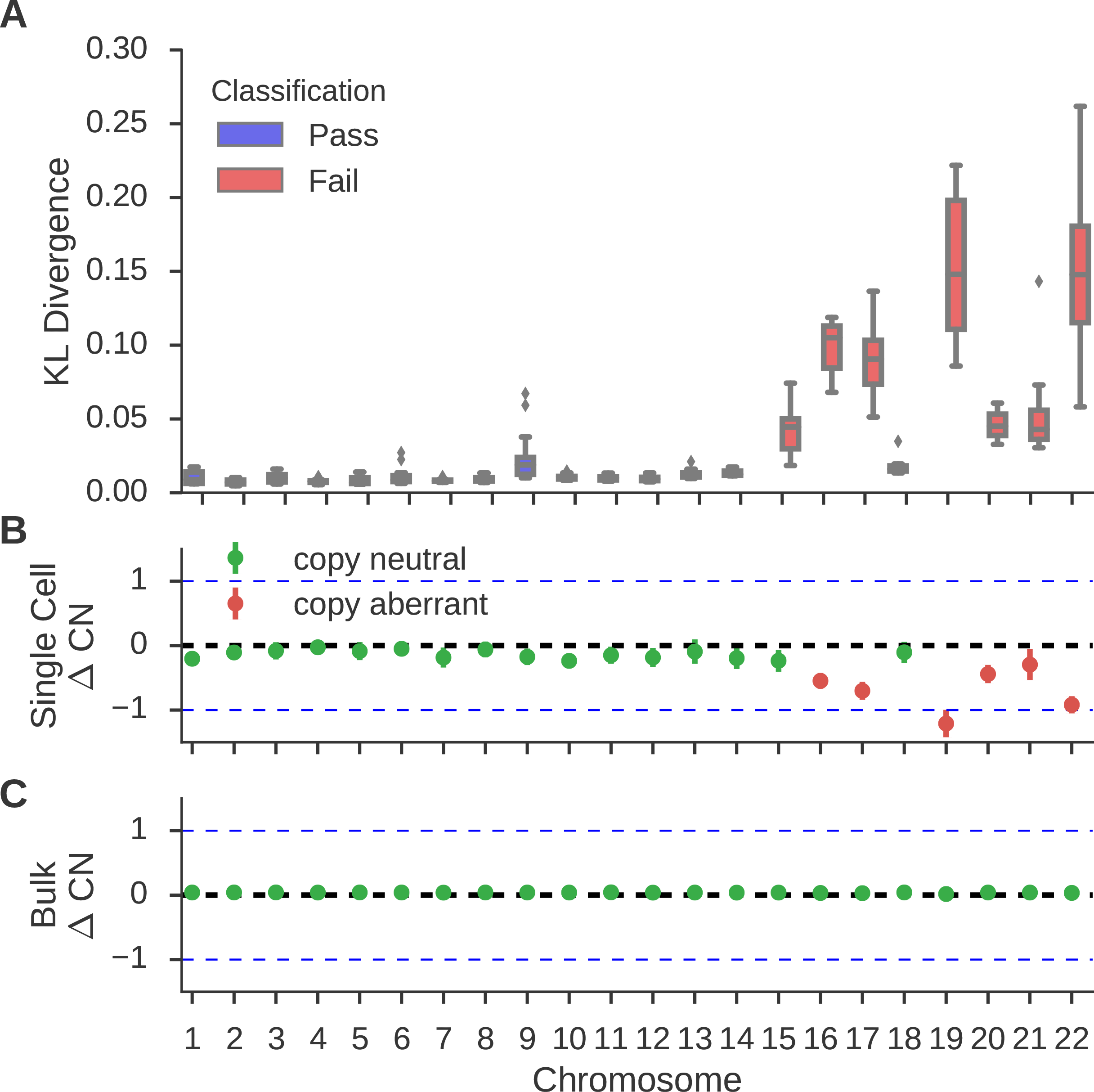
Identification of false-positive chromosomal copy changes due to poor amplification. **A**. boxplots of the KL divergence of each autosome from the sample-average PSD for the 16 “1465” samples. Chromosomes are labeled as failed (red) if PaSD-qc identifies the chromosome as aberrantly amplified in at least half of the samples (Table S2). **B**. the average copy number across all samples as inferred by the BICseq2 algorithm. Errorbars represent standard deviation across all samples. Chromosomes are considered copy aberrant if BICseq2 identifies a significant (p-value < 0.05) alteration in at least 25% of samples (Table S2). A chromosomal deletion in at least 25% of cells should be identifiable in bulk sequencing. **C**. chromosome copy profile of a bulk sample from the same tissue as the single cell samples. All chromosomes are copy neutral.

### Discriminating high- and low-quality samples

We additionally profiled three newly amplified samples from the “1465” individual. Prior analysis showed these samples to be of low quality (Figure S5). Comparing them to high-quality samples from “1465” and “4638” provide an illustrative example of how PaSD-qc distinguishes high- and low-quality samples. Not only are the PSDs distinguishable by eye (Figure 6A), but the poor-quality samples also have a wider distribution of amplicon sizes and smaller median amplicon size (Figure 6B). Additionally, using the symmetric KL-divergence, PaSD-qc clusters the libraries based on amplification behavior (Figure 6C). The clustering correctly groups samples by high- and low-quality and further by biological origin. Finally, PaSD-qc can use the symmetric KL-divergence to probabilistically assign samples to different categories (e.g., high- and low-quality) using pre-computed gold-standard spectra. The toolbox includes methods which allow users to generate these gold-standard spectra from their own data. PaSD-qc can fully and accurately profile samples with coverage as low as 0.5X, and it provides accurate sample clustering and category assignment with coverage as low as 0.1X (Figure S6).

**Figure 6:**
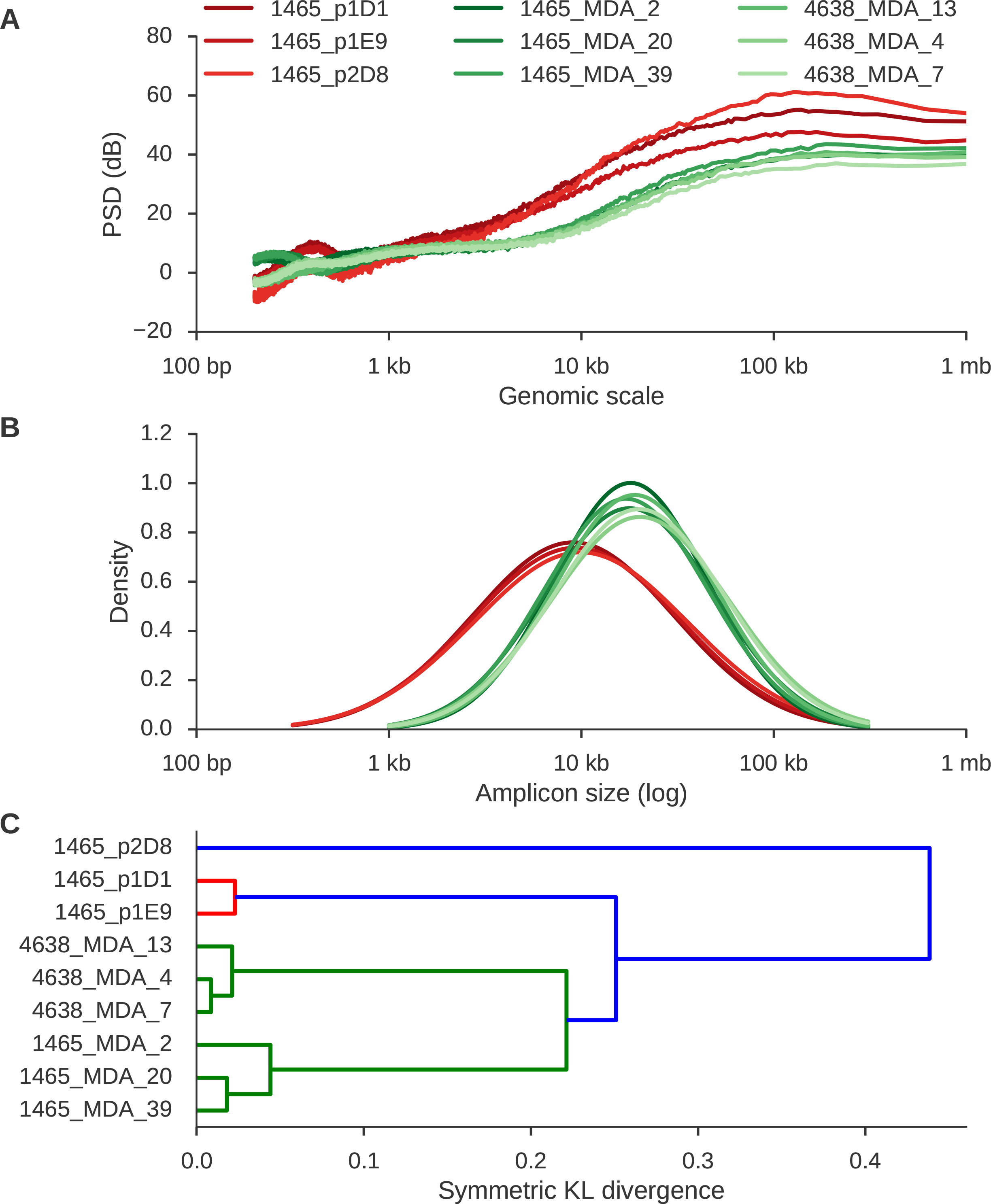
PaSD-qc separates high-quality from low-quality samples and groups similarly behaving libraries. **A**. power spectral densities for three low-quality (red) and six high-quality libraries (green). **B**. Amplicon size density plots for the nine samples. **C**. Hierarchical clustering using the symmetric KL-divergence correctly groups the samples based on both quality and biological origin.

## Discussion

Here we have demonstrated the effectiveness of PaSD-qc to comprehensively evaluate the quality and amplification properties of scWGS data. Although several studies have recently compared the uniformity of different scWGS protocols [21, 24, 25], each study uses its own collection of statistics, making the task of determining the superior protocol difficult. We believe PaSD-qc represents an important step forward for the field as it provides a standardized suite of analyses that researchers can easily insert into any pipeline. In particular, PaSD-qc introduces novel methods to estimate the full distribution of amplicon sizes in a sample, identify individual chromosomes which were poorly amplified, and compare samples based on amplification behavior.

These analyses not only allow comparisons across amplification protocols but also provide an important starting point for variant analysis. For example, it was recently demonstrated that the correlation in allelic balance induced by the large amplicons of MDA can be exploited to increase the accuracy of single cell single nucleotide variant (SNV) calling [11]. Dong et al. proposed a method employing a kernel smoothing algorithm that requires a user-defined bandwidth to compute the expected balance at a given genomic locus. The length of the bandwidth reflects the user's belief about the maximum distance at which informative correlation exists, and the authors suggest using a fixed bandwidth of 10 kb for all samples. However, PaSD-qc provides a principled, data-driven strategy to assign a tailored bandwidth to each individual sample as the 95% upper bound on amplicon sizes naturally defines a maximum correlation distance.

Additionally, our results address the question of whether *in vitro* amplification of the human genome by the Phi-29 MDA polymerase [26] produces amplicons of 10-100 kb as documented in bacterial genomes [27]. We found that some protocols approach the upper bound while others produce far smaller amplicons, with the lower bound in the 1-5 kb range. This has important consequences for PacBio or 10X Genomics sequencing on single cells in which fragments many kilobases in length are required. In particular, only some protocols may consistently produce large enough amplicons to make long-insert or haplotype-based sequencing possible.

Our results demonstrate that single cell amplification methods can artifactually induce whole chromosome copy alterations due to systematic under-amplification. Patterns of under amplification appear to be consistent across the same protocol but to vary between different protocols. As scWGS is becoming an increasingly popular choice to characterize copy number alterations in both research and clinical settings [6, 16, 20], the ability to identify false-positive copy changes is important. In addition, our results suggest that high-quality MDA data are likely yield accurate calls for small CNVs (smaller than its amplicon sizes), as its sub-amplicon variance approaches that of bulk sequencing. Prior studies have focused on the detection of large copy alterations; none have specifically examined suitability for very small CNV calling.

Lastly, full mutational analysis at the single cell level requires high-coverage (>30X) sequencing, but the uneven quality of scWGS data, primarily due to the variable quality of cells, has often resulted in only a portion of the data generated being usable. The ability to accurately characterize data quality from low-coverage data suggests that a cost-effective approach in scWGS data generation is to screen a large number of cells at very low coverage (e.g., <0.1X) and select only a small number of high-quality candidates for additional sequencing. PaSD-qc provides an efficient computational framework to perform this evaluation.

## Conclusion

High-coverage scWGS enables identification of single nucleotide variants and other mutations at the single cell level, but mitigating the biases arising from whole-genome amplification remains a challenge. The proposed statistical method allows a detailed characterization of the data quality for scWGS datasets, aiding in selection of appropriate protocols and ensuring the fidelity of downstream analysis.

## Methods

### Data

MDA data for “4638” (Brain A), “1465” (Brain B), and “4643” (Brain C) were previously obtained by our group [4]. Additionally, three muscles cells from the “1465” individual were isolated, amplified, and sequenced as in that study. Fourteen additional MDA samples (C1a/b, C2a/b, C3a, N1a/b, N2a/b, N3a, N4a/b) were obtained from [20] (Short Read Archive accession number SRP052954). MALBAC samples were obtained from [5]. In Figure 2, the bulk sample is bulk cortex from “1465”, the MDA sample is cell 30 from “1465”, and the MALBAC sample has the SRA accession number SRX204745. All data were downsampled to 1X using SAMtools prior to analysis.

### Power spectral density estimation

Starting with a BAM file, read depth for each arm of each chromosome is extracted as the time series 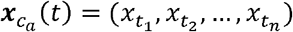 where *c* is the chromosome, *a* is the chromosome arm and *t_i_* is the start position of a uniquely mappable read. Uniquely mappable positions for the hg19 genome were download from the UCSC genome browser. By default, PaSD-qc uses mappability tracks calculated for 100 bp reads. Any series with fewer than 10 million observations is removed from further analysis. Each series 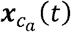 is then divided into *M* windows of length *L* overlapping by *D* positions. By default, *L* = 1×10^6^ and D = 5×10^5^. The Lomb-Scargle algorithm [17, 18] is used to calculate the power spectral density, 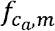, for each 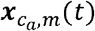 at eight thousand frequencies, *ω*, evenly spaced from 1e-6 to 5e-3. The PSD for each chromosome is then estimated as

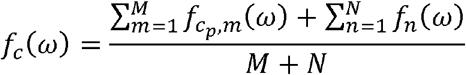

where *M* and *N* are the number of windows on the *p* and *q* arms of chromosome *c*, respectively. The average PSD for an individual sample is then calculated as *f*(*ω*) = median{*f_c_*(*ω*)}.

The mathematical details of Lomb-Scargle PSD estimation are described in SI. The theoretical justification for the power spectral density as a measure of variance in an aperiodic signal is also given in SI.

### Normalizing and plotting power spectral densities

To remove edge effects and effects arising purely from sequencing, we take an idealized bulk sample as the baseline for the read depth power spectral density. The idealized bulk PSD, *f_b_*, was derived by fitting a lowess curve to the bulk PSD shown in Figure 2A. The spectral density for each single cell sample is then normalized using the decibel transform as

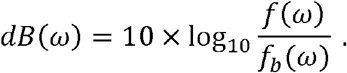

This transform is standard in digital signal processing to remove a background signal.

Traditionally, power spectral densities are plotted as a function of frequency. However, for the genomic read depth signal, frequency takes on the unintuitive units of inverse genomic scale (1/bp). We instead choose to plot the PSD as a function of period, 1/ω. This results in the familiar units of genomic scale (bp) on the x-axis. We believe this eases interpretation, especially for those unfamiliar with power spectral densities.

### Estimating the distribution of amplicon sizes from the power spectral density

As motivated in “Results”, the dynamic portion of the scWGS PSD curve reflects the cumulative distribution of the amplicon sizes in that sample. This distribution can thus be estimated by fitting a linearly scaled cumulative distribution function to this dynamic region. In practice, which distribution function should be fit is governed by two principles: 1) how tractable is fitting the curve using modern gradient descent algorithms, and 2) how well does the estimated distribution reproduce the original data. The first problem is one purely of computation and amounts to whether the distribution function has a closed-form solution or easily approximated integral solution. We tested three distributions which fit this criterion: the normal (erf), logistic, and gamma distributions. To solve the second problem, we used the estimated density to simulate an idealized amplification process and compared the PSD of the idealized process to that of the original sample. The simulation procedure is described in the section below. We found the normal (erf) distribution best reproduced the data.

Let 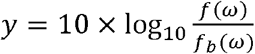 and *x* = −log_10_*ω*. The dynamic region of the curve is fit as

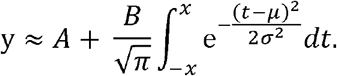

The log-transformed density of the amplicon sizes is then estimated as *𝒩(μ,σ^2^)*. To estimate the median and 95% bounds, we draw 100,000 observations from the above distribution and calculate the median and percentiles of 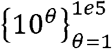, where *θ* is a simulated observation.

### Simulating an idealized amplification process

Let *p*(·|Θ) be the log-distribution of amplicon sizes estimated using the above method. For a given chromosome arm, an idealized amplification process is simulated using the following algorithm:

1. Initialize a vector, *v*, of length equal to the length of chromosome arm with all entries zero.
2. Randomly simulate an amplicon size as *l* = 10^*θ*^ where *θ*~*p*(·|Θ).
3. Randomly choose a starting position *s*, where *s*~Unif(*a,b*) where *a* and *b* are the start and end coordinates of the chromosome arm
4. Increase the values of the entries of *v* overlapped by the amplicon by one

a. Note: if *s* + *l* > *b*, the simulated amplicon is discarded
5. Repeat 2-5 until the desired average depth of coverage is reached.

a. Depth of coverage is calculated as 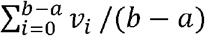.
6. Randomly choose *K* non-zero observations from *v* where *K* is the number of non-zero observations from the chromosome arm in the original data.

The PSD of the resulting simulated read depth signal is then estimated and normalized as described above. To account for total power differences and mean shifts between the simulated data and the true data due to the idealized nature of the above algorithm, we normalize each curve by the maximum observed power and mean shift each curve such that *f*(10^−3^) = 0. We chose to use the p arm of chromosome 3 for simulation purposes as in our experience it is a large arm with highly consistent amplification across samples.

### Estimating the autocovariance function

The autocovariance function, *γ*, estimates the covariance of a time series against itself at lags *k*. As derived in SI, the real-valued sample autocovariance can be estimated from the PSD as

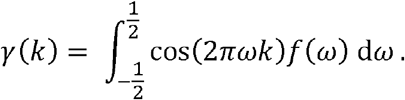

This integral can be quickly and accurately estimated numerically using any modern quadrature technique. We use Simpson's rule.

To directly calculate the ACF from unevenly space time series data, we define the “observation” function as

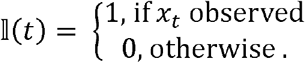

For lag *h* we construct the set *S_h_* = {*x_t_* | 𝕀 (*t* + *h*) = 1}, which is the set of all observations such that an observation at a distance of *h* is also present. The sample autocovariance is then calculated as

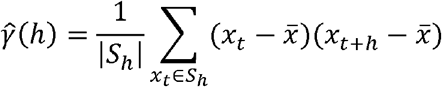

where *x̄* is the sample mean of the time series and ∣*s_h_*∣ denotes the size of *s_h_*.

### Comparing the behavior of different spectra

Given two probability densities, *p*_1_ and *p*_2_ and a vector of observations, ***X***, the Kullback-Leibler divergence is an informatic dissimilarity measure between the two densities and is defined as

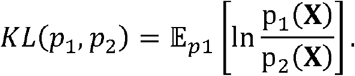

It can be shown (see SI) that the Kullback-Leibler (KL) divergence between two PSDs is

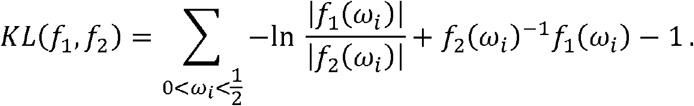

The KL-divergence is not a true distance metric as *KL*(*f*_1_,*f*_2_) ≠ *KL*(*f*_2_,*f*_1_). Following [22], we define the symmetric divergence between to spectra as

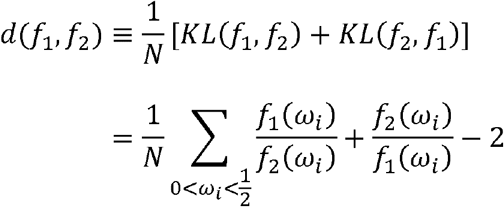

where *N* is the total number of frequencies in the sum. This value is reflexive and always non-negative (see SI); thus *d* is a principled statistical distance metric between two spectra.

To identify aberrantly amplified chromosomes, we calculate *d*(*f, f_c_*) for each chromosome of a sample. We then calculate the median divergence and the median absolute difference of the divergences. A chromosome is considered aberrant if its divergence is greater than the sum of the median and two times the median absolute difference. To cluster samples by behavior, the pairwise divergence is calculated between each pair of sample PSDs. The resulting symmetric distance matrix is then used to perform hierarchical clustering.

### Estimating median absolute pairwise difference

The BICseq2 algorithm [23] was used to calculate the copy number in bins of 1 kb, 5kb, 10 kb, 50 kb, 100 kb, 500 kb, and 1 mb for all “1465” samples. Estimates were corrected for mappability and GC content. For each bin size, MAPD is calculated as median 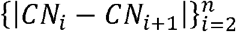, where *CN_i_* is the copy number in the i^th^ bin and *n* is the total number of bins.

### Estimating chromosome-level copy number

The BICseq2 algorithm was used to calculate the copy number in bins of 500 kb normalized for mappability and GC content. The copy number for each chromosome was taken to be the median copy number over all bins overlapping that chromosome. Additionally, BICseq2 automatically assigns a p-value to the significance of the copy change, and a chromosome was considered significantly copy altered if the assigned p-value was less than 0.05.

### Implementation

PaSD-qc is implemented in python. It uses SAMtools to extract coverage from bam files and the astropy package [28] to implement an *O*(*n* · log *n*) Lomb-Scargle algorithm. The function curve_fit in the scipy module is used to fit the modified erf function to the scWGS PSD. Clustering of samples is performed by the linkage function also in the scipy module. PaSD-qc parallelizes across samples for efficient multi-sample analysis. Source code, documentation, and examples – including all data and code to reproduce the figures in this manuscript – are available at https://github.com/parklab/PaSD-qc.

## Abbreviations

scWGS: single cell whole-genome sequencing
PSD: power spectral density
ACF: autocovariance function
MDA: multiple displacement amplification
MALBAC: multiple looping and annealing based amplification cycles
KL-divergence: Kullback-Leibler divergence
CNV: copy number variation
SNV: single nucleotide variation
Erf: error function

## Acknowledgements

We thank Joe Luquette, Doga Gulhan, and Alon Galor for their valuable input in preparing this manuscript. We thank the Brain Somatic Mosaicism Network for supporting this research.

## Declarations

### Availability of data and materials

PaSD-qc is implemented as a python package which is freely available at https://github.com/parklab/PaSD-qc along with all data and scripts necessary to reproduce the figures in this paper. BAM files were downloaded from the Short Read Archive with accession numbers SRP042470 for “1465” from [4], SRP061939 for “4638” and “4643” from [4], SRA060929 for samples from [5], and SRP052954 for samples from [20]. The BAMs for the three additional “1465” samples shown in Figure 6 are available from the corresponding author upon reasonable request.

### Author's contributions

MAS and PJP conceived the idea for the paper. MAS designed the statistical methods, implemented the algorithms, performed the analysis, and wrote the manuscript. ARB and CV aligned the samples and helped prepare the manuscript. ML and MEC collected the data. CAW provided resource support and read the manuscript.

### Consent for publication

Not applicable.

### Ethics approval and consent to participate

Not applicable

### Additional files

Additional file 1: supplementary information, tables S1-S2, and figures S1-S6

## References

[1] G. D. Evrony, E. Lee, B. K. Mehta, Y. Benjamini, R. M. Johnson, X. Cai, L. Yang, P. Haseley, H. S. Lehmann, P. J. Park, and C. A. Walsh, “Cell lineage analysis in human brain using endogenous retroelements.” Neuron, vol. 85, pp. 49–59, Jan. 2015.

[2] Y. Wang, J. Waters, M. L. Leung, A. Unruh, W. Roh, X. Shi, K. Chen, P. Scheet, S. Vattathil, H. Liang, A. Multani, H. Zhang, R. Zhao, F. Michor, F. Meric-Bernstam, and N. E. Navin, “Clonal evolution in breast cancer revealed by single nucleus genome sequencing.” Nature, vol. 512, pp. 155–160, Aug. 2014.

[3] M. J. McConnell, M. R. Lindberg, K. J. Brennand, J. C. Piper, T. Voet, C. Cowing-Zitron, S. Shumilina, R. S. Lasken, J. R. Vermeesch, I. M. Hall, and F. H. Gage, “Mosaic copy number variation in human neurons.” Science, vol. 342, pp. 632–637, Nov. 2013.

[4] M. A. Lodato, M. B. Woodworth, S. Lee, G. D. Evrony, B. K. Mehta, A. Karger, S. Lee, T. W. Chittenden, A. M. D'Gama, X. Cai, L. J. Luquette, E. Lee, P. J. Park, and C. A. Walsh, “Somatic mutation in single human neurons tracks developmental and transcriptional history.” Science, vol. 350, pp. 94–98, Oct. 2015.

[5] C. Zong, S. Lu, A. R. Chapman, and X. S. Xie, “Genome-wide detection of single-nucleotide and copy-number variations of a single human cell.” Science, vol. 388, pp. 1622–1626, December 2012.

[6] W. Liu, H. Zhang, D. Hu, S. Lu, and X. Sun, “The performance of MALBAC and MDA methods in the identification of concurrent mutations and aneuploidy screening to diagnose beta-thalassaemia disorders at the single- and multiple-cell levels.” Journal of clinical laboratory analysis, May 2017.

[7] J. Xu, R. Fang, L. Chen, D. Chen, J.-P. Xiao, W. Yang, H. Wang, X. Song, T. Ma, S. Bo, C. Shi, J. Ren, L. Huang, L.-Y. Cai, B. Yao, X. S. Xie, and S. Lu, “Noninvasive chromosome screening of human embryos by genome sequencing of embryo culture medium for in vitro fertilization.” Proceedings of the National Academy of Sciences of the United States of America, vol. 113, pp. 11907–11912, Oct. 2016.

[8] T. Baslan, J. Kendall, L. Rodgers, H. Cox, M. Riggs, A. Stepansky, J. Troge, K. Ravi, D. Esposito, B. Lakshmi, M. Wigler, N. Navin, and J. Hicks, “Genome-wide copy number analysis of single cells.” Nature protocols, vol. 7, pp. 1024–1041, May 2012.

[9] Y. Wang and N. E. Navin, “Advances and applications of single-cell sequencing technologies.” Molecular cell, vol. 58, pp. 598–609, May 2015.

[10] C.-Z. Zhang, V. A. Adalsteinsson, J. Francis, H. Cornils, J. Jung, C. Maire, K. L. Ligon, M. Meyerson, and J. C. Love, “Calibrating genomic and allelic coverage bias in single-cell sequencing.” Nature communications, vol. 6, p. 6822, Apr. 2015.

[11] X. Dong, L. Zhang, B. Milholland, M. Lee, A. Y. Maslov, T. Wang, and J. Vijg, “Accurate identification of single-nucleotide variants in whole-genome-amplified single cells.” Nature methods, vol. 14, pp. 491–493, May 2017.

[12] M. L. Leung, Y. Wang, J. Waters, and N. E. Navin, “SNES: single nucleus exome sequencing.” Genome biology, vol. 16, p. 55, Mar. 2015.

[13] K. Leung, A. Klaus, B. K. Lin, E. Laks, J. Biele, D. Lai, A. Bashashati, Y.-F. Huang, R. Aniba, M. Moksa, A. Steif, A.-M. Mes-Masson, M. Hirst, S. P. Shah, S. Aparicio, and C. L. Hansen, “Robust high-performance nanoliter-volume single-cell multiple displacement amplification on planar substrates.” Proceedings of the National Academy of Sciences of the United States of America, vol. 113, pp. 8484–8489, July 2016.

[14] M. Rhee, Y. K. Light, R. J. Meagher, and A. K. Singh, “Digital droplet multiple displacement amplification (ddMDA) for whole genome sequencing of limited DNA samples.” PloS one, vol. 11, 2016.

[15] C. Chen, D. Xing, L. Tan, H. Li, G. Zhou, L. Huang, and X. S. Xie, “Single-cell whole-genome analyses by linear amplification via transposon insertion (LIANTI).” Science, vol. 356, pp. 189–194, Apr. 2017.

[16] X. Cai, G. D. Evrony, H. S. Lehmann, P. C. Elhosary, B. K. Mehta, A. Poduri, and C. A. Walsh, “Single-cell, genome-wide sequencing identifies clonal somatic copy-number variation in the human brain.” Cell reports, vol. 8, pp. 1280–1289, Sept. 2014.

[17] N. R. Lomb, “Least-squares frequency analysis of unequally spaced data.” Astrophysics and Space Science, vol. 39, pp. 447–462, Feb. 1976.

[18] J. D. Scargle, “Studies in astronomical time series analysis. ii - statistical aspects of spectral analysis of unevenly spaced data.” Astrophysical Journal, vol. 263, pp. 835–853, Dec. 1982.

[19] P. Welch, “The use of fast Fourier transform for the estimation of power spectra: A method based on time averaging over short, modified periodograms.” IEEE Transactions on Audio and Electroacoustics, vol. 15, pp. 70–73, June 1967.

[20] C.-Z. Zhang, A. Spektor, H. Cornils, J. M. Francis, E. K. Jackson, S. Liu, M. Meyerson, and D. Pellman, “Chromothripsis from DNA damage in micronuclei.” Nature, vol. 522, pp. 179–184, June 2015.

[21] L. Ning, Z. Li, G. Wang, W. Hu, Q. Hou, Y. Tong, M. Zhang, Y. Chen, L. Qin, X. Chen, H.-Y. Man, P. Liu, and J. He, “Quantitative assessment of single-cell whole genome amplification methods for detecting copy number variation using hippocampal neurons.” Scientific reports, vol. 5, p. 11415, June 2015.

[22] R. H. Shumway and D. S. Stoffer, “Statistical methods in the frequency domain,” in Time Series Analysis and Its Applications, Springer, Jan. 2011.

[23] R. Xi, S. Lee, Y. Xia, T.-M. Kim, and P. J. Park, “Copy number analysis of whole-genome data using bic-seq2 and its application to detection of cancer susceptibility variants.” Nucleic acids research, vol. 44, pp. 6274–6286, July 2016.

[24] C. F. A. de Bourcy, I. De Vlaminck, J. N. Kanbar, J. Wang, C. Gawad, and S. R. Quake, “A quantitative comparison of single-cell whole genome amplification methods.” PloS one, vol. 9, 2014.

[25] E. Borgström, M. Paterlini, J. E. Mold, J. Frisen, and J. Lundeberg, “Comparison of whole genome amplification techniques for human single cell exome sequencing.” PloS one, vol. 12, 2017.

[26] F. B. Dean, S. Hosono, L. Fang, X. Wu, A. F. Faruqi, P. Bray-Ward, Z. Sun, Q. Zong, Y. Du, J. Du, M. Driscoll, W. Song, S. F. Kingsmore, M. Egholm, and R. S. Lasken, “Comprehensive human genome amplification using multiple displacement amplification.” Proceedings of the National Academy of Sciences of the United States of America, vol. 99, pp. 5261–5266, Apr. 2002.

[27] L. Blanco, A. Bernad, J. M. Lázaro, G. Martín, C. Garmendia, and M. Salas, “Highly efficient DNA synthesis by the phage phi 29 DNA polymerase. symmetrical mode of DNA replication.” The Journal of biological chemistry, vol. 264, pp. 8935–8940, May 1989.

[28] A. Collaboration, T. P. Robitaille, E. J. Tollerud, P. Greenfield, M. Droettboom, E. Bray, T. Aldcroft, M. Davis, A. Ginsburg, A. M. Price-Whelan, W. E. Kerzendorf, A. Conley, N. Crighton, K. Barbary, D. Muna, H. Ferguson, F. Grollier, M. M. Parikh, P. H. Nair, H. M. Unther, C. Deil, J. Woillez, S. Conseil, R. Kramer, J. E. H. Turner, L. Singer, R. Fox, B. A. Weaver, V. Zabalza, Z. I. Edwards, K. Azalee Bostroem, D. J. Burke, A. R. Casey, S. M. Crawford, N. Dencheva, J. Ely, T. Jenness, K. Labrie, P. L. Lim, F. Pierfederici, A. Pontzen, A. Ptak, B. Refsdal, M. Servillat, and O. Streicher, “Astropy: A community python package for astronomy,”, vol. 558, p. A33, Oct. 2013.

